# Revealing Interactions between Microbes, Metabolites, and Dietary Compounds using Genome-scale Analysis

**DOI:** 10.64898/2025.12.03.692129

**Authors:** Tong Wang, Benjamin Gyori, Scott T. Weiss, Giulia Menichetti, Yang-Yu Liu

## Abstract

**Background:** The role of gut microbiome in predicting diet response and developing personalized dietary recommendations has been increasingly recognized. Yet, we still lack comprehensive, genome-based insights into which gut microbes metabolize specific dietary compounds.

**Results:** Here, we leveraged the metabolic networks constructed from well-annotated microbial genomes to characterize the potential interactions between microbes and metabolites, specifically emphasizing the interactions between microbes and dietary compounds. We revealed a substantial, approximately four-fold variation in both the number of metabolites and dietary compounds in the microbial genome-scale metabolic networks across different genera, whereas species within the same genus showed a high metabolic similarity (mean coefficient of variation in microbial network degree *CV* = 0.023 for metabolites and 0.015 for dietary compounds). We found that the number of species that can utilize a metabolite drastically varies, ranging from 1 to 818 species, with some metabolites being used by a wide range of species (211 out of 1390 metabolites used by more than 95% of species) and others only by a few species (435 metabolites used by less than 5% of species). Leveraging a longitudinal microbiome study, we observed that microbial taxa with similar metabolic capacity tend to have positively correlated abundances, and the gut microbiome’s capacity to process dietary compounds is functionally stable. Finally, we propose a network-based method to identify the dietary compounds that are specific to no more than 10 microbial species, offering a new strategy for combining a dietary compound and its linked microbial species to design synbiotics.

**Conclusions:** Our results quantitatively reveal large-scale variation and redundancy in gut microbial metabolism and identify dietary compounds linked to only a few microbial species. These findings improve understanding of microbe-metabolite interactions and provide a foundation for the rational design of microbiome-based interventions for healthy benefits.

## Introduction

The gut microbiome significantly influences host physiological functions, with one of the primary mechanisms being the consumption of dietary molecules and the production of metabolic byproducts^1–3^. For instance, non-digestible carbohydrates^1,2,4,5^ and flavonoids^6,7^, which humans cannot digest, act as a vital energy source for certain gut microbes. As a result, these microbes produce important metabolites, such as short-chain fatty acids (SCFAs) and essential amino acids, which can affect the host’s metabolism and immune function and are essential for maintaining overall health^1,8^. Therefore, to design dietary or microbiome-targeted interventions, it is essential to extensively map the complex interactions among microbes, dietary compounds, and metabolites^9–14^.

Over the past decade, substantial progress has been made toward linking diet, microbial community structure, and metabolic function in both gnotobiotic mice and human cohorts^15–20^. More recently, shotgun metagenomics and integrative multi-omics approaches have provided higher-resolution insights into microbial functional capacity, revealing how dietary factors shape these functions and uncovering inter-species variation in nutrient utilization pathways^15,17,19^. These efforts have collectively advanced our understanding of metabolic specialization and dietary preference within the gut microbiome.

However, despite these advances, a comprehensive genome-scale assessment of complex food-microbe-metabolite relationships remains limited^21,22^. Most studies have focused on a few well-characterized taxa (e.g., *Eubacterium ramulus*^6,7^, *Ruminococcus torques*^18^) or specific compound classes (e.g., carbohydrates^1,2,4,5^ and flavonoids^6,7^), leaving broader taxonomic and metabolic diversity underexplored. Adopting a more comprehensive network perspective will enhance our understanding of the gut microbiome’s functional redundancy (i.e., the extent to which multiple species contribute to the same function) and its resilience to dietary changes, bridging significant knowledge gaps in microbe-metabolite interactions and informing precise dietary or microbiome-targeted interventions.

Here, to fill this knowledge gap, we leveraged the microbial genomes of representative gut microbes available in AGREDA (AGORA-based REconstruction for Diet Analysis)^23,24^, a comprehensive database comprising genome-scale metabolic models (GEMs) for over 800 microbial species with curated biochemical annotations that enable systematic mapping of potential microbe–metabolite interactions. Built upon AGORA (Assembly of Gut Organisms through Reconstruction and Analysis)^25,26^, AGREDA incorporates additional dietary compounds as new nodes in metabolic networks encoded by the microbial genomes^23,24^. By analyzing these networks, we explored the interactions between microbes, metabolites, and dietary compounds, ecological interactions among microbes, and distinct ecological and network patterns of dietary compounds compared to metabolites. Our findings provide a systems-level understanding of the metabolic diversity, redundancy, and potential for rational design of synbiotics (combinations of beneficial microorganisms and dietary supplements) in the human gut microbiome.

## Results

### Inference of microbe-metabolite interactions

While some databases include experimentally verified interactions^27,28^, they are limited in coverage^29^. To expand this, we used metabolic networks built from microbial genomes in the AGREDA database^23,24^. AGREDA extends AGORA^25,26^ by adding more dietary metabolism pathways (see **Methods** for details). As a result, AGREDA has 312 dietary compounds, more than 99 of which exist in AGORA.

In this study, metabolites refer to the complete set of all chemicals represented in the metabolic networks of all microbial species in AGREDA. Dietary compounds are a subset of these metabolites that are derived from the human diet and documented in the i-Diet database (http://www.i-diet.es), a comprehensive repository containing 650 dietary compounds. To understand the statistical properties of the metabolites represented in AGREDA, we leveraged ClassyFire^30^ and found that the five chemical classes of dietary compounds are flavonoids, organooxygen compounds, fatty acyls, prenol lipids, and phenols.

For each microbial genome in AGREDA, we constructed its corresponding metabolic network and documented all existing metabolites within that network (**Fig. 1a**). We then created an incidence matrix of microbe-metabolite interactions, where each element indicates the presence or absence of a metabolite in a species’ metabolic network (**Fig. 1a**). This approach allowed us to leverage microbial genomes to infer potential interactions between microbes and metabolites, encompassing a total of 818 microbial species and 1390 metabolites. For the 818 microbial genomes in AGREDA, the median fraction of annotated genes among all coding genes is 97.6%, which verifies the high quality of the microbial genomes in AGREDA.

**Figure 1:**
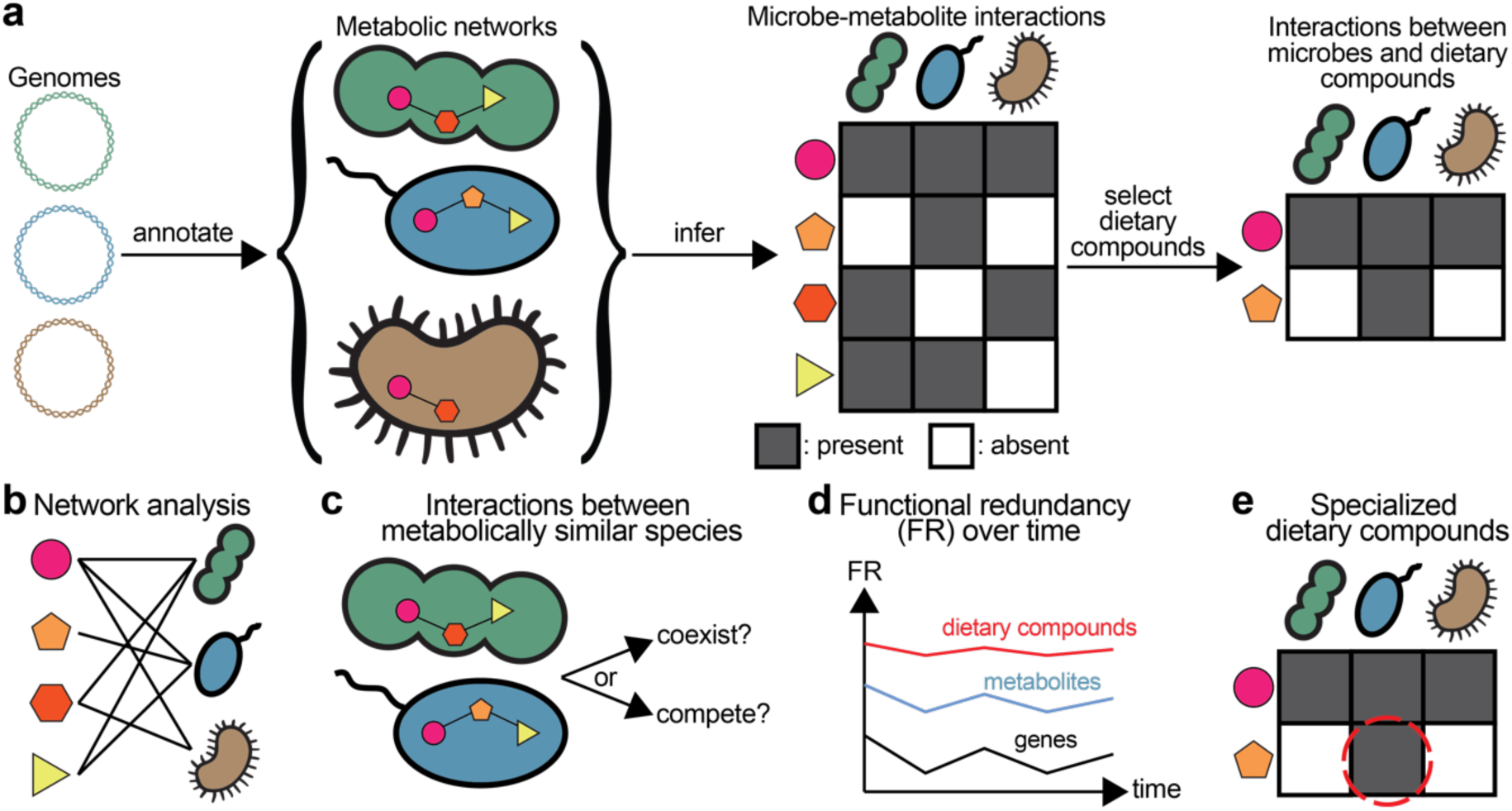
Genome-based inference of microbe-metabolite interactions and their characterization. **a**, Inferring microbe-metabolite interactions based on microbial genomes. The microbial genome of a species is annotated to obtain its metabolic network, comprising many metabolites linked by chemical reactions. Based on the metabolic network, we created the incidence matrix for microbe-metabolite interactions, who’s each element denotes whether a metabolite exists in the metabolic network of one species. The grey/white color of each component represents the presence/absence of a metabolite in the metabolic network of a species. The interactions between microbes and dietary compounds can be constructed by selecting dietary compounds from metabolites. **b,** Network analysis of the microbe-metabolite interactions. **c,** The microbe-metabolite interactions can be used to infer the interactions between metabolically similar species. **d,** The constructed interactions can be used to compute the functional redundancy for dietary compounds, metabolites, or genes. **e,** The dietary compounds that are specialized by a few species can be used to target the microbes that can utilize them.

### Analysis of microbe-metabolite interactions

We represented microbe-metabolite interactions as a bipartite graph (**Fig. 2a**), where a microbial species is linked to a metabolite if that metabolite plays a role in its metabolic network. We grouped microbial species by their genera and metabolites by their chemical classes in ClassyFire^30^. The edge density of the bipartite graph (i.e., the fraction of edges present in the bipartite network out of the total possible edges between microbial species and metabolites) is 39.84%. By contrast, when examining other manually curated databases based on experimentally tested yet unreported microbe-metabolite interactions, we observed a much lower link density, e.g., 3.63% for NJS16^27^ and 3.38% for NJC19^28^. This indicates that manually curated databases^29^ are subject to underreporting, rendering them unsuitable for our systematic analysis.

**Figure 2:**
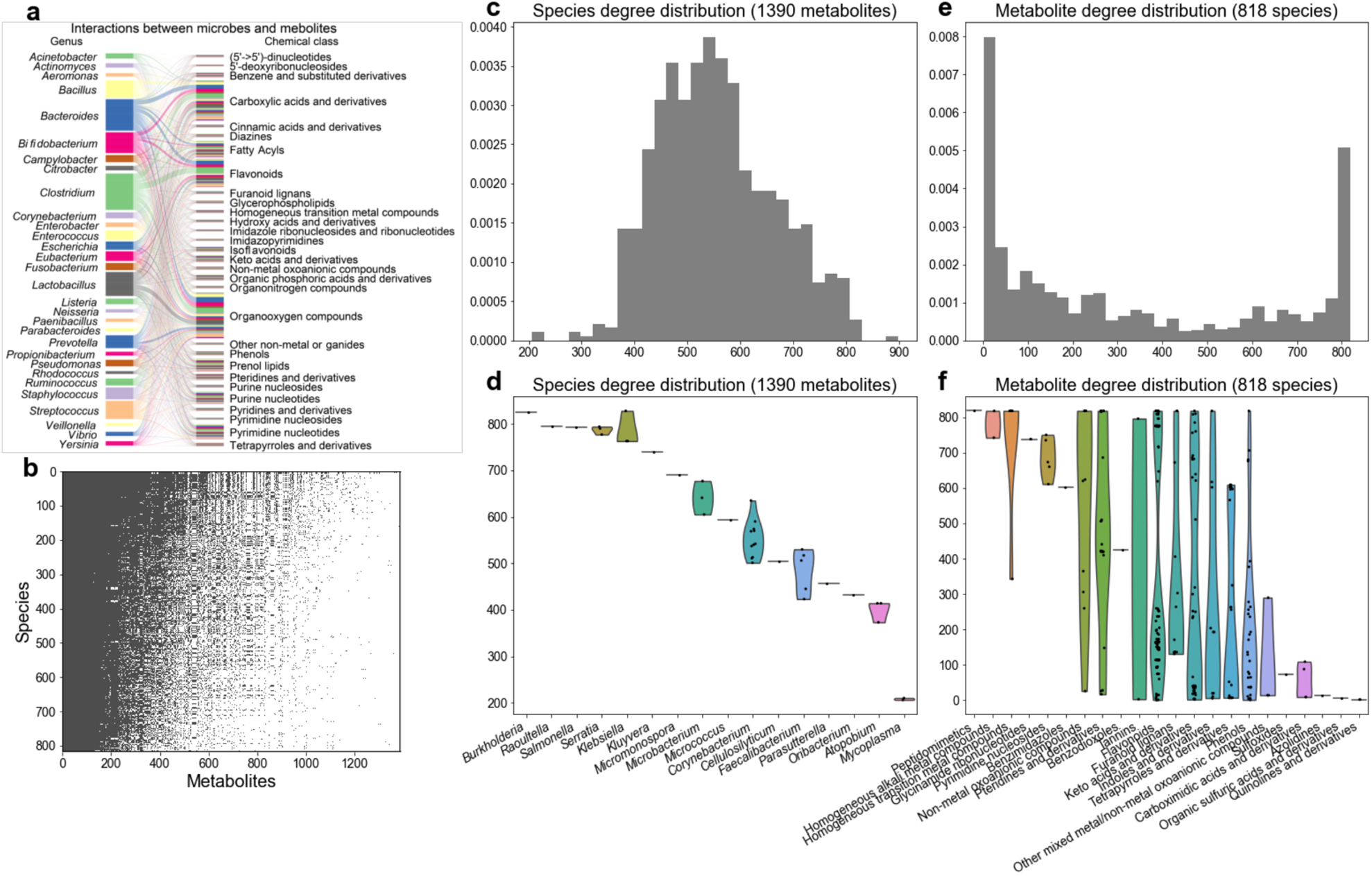
The inferred interactions between microbes and metabolites. **a,** The bipartite network of interactions. To avoid visual overcrowding, we only display the top 30 genera and chemical classes with the largest counts. Each thin line on the left or right side corresponds to one species or metabolite, respectively. Different species from the same genus and different metabolites from the same chemical class are grouped together without any gap in between. The curved line in the middle connects a species with a metabolite when they interact. The color of nodes and edges is assigned based on the genus on the left. **b,** The incidence matrix of the interactions. The grey/white color for each element represents the presence/absence of a metabolite in the metabolic network of a species. This matrix is organized using the Nestedness Temperature Calculator^71^ to emphasize its nested structure. **c,** A histogram of the species degree distribution with the number of bins chosen as 30. **d,** The species degree distribution for species from representative genera. **e,** A histogram of the metabolite degree distribution with the number of bins chosen as 30. **f,** The metabolite degree distribution from representative chemical classes.

By analyzing the incidence matrix of the microbe-metabolite bipartite graph (**Fig. 2b**), we observed a highly nested structure (with nestedness metric NODF = 0.704). This indicates that for those species with lower metabolic capacity (i.e., interacting with fewer metabolites), their metabolites tend to be a subset of the metabolites of those species with higher metabolic capacity. This result is consistent with our previous finding that the gene content network of microbial species also has a highly nested structure, meaning that for those species with smaller genomes, their genes tend to be part of those species with larger genomes^31^. From an evolutionary standpoint, nestedness is intriguing because it reflects underlying processes such as species co evolution^32^ and gene sharing through horizontal gene transfer^31^ (see **Methods**).

To reveal the difference in the metabolic capacity across different microbial species, we calculated the species degree distribution (with ‘degree’ the number of metabolites that each species is connected with), which appears mean-centered and unimodal (**Fig. 2c**). We then compared the metabolic capacity of species in different genera, finding that species from the same genus have similar metabolic capacity (mean coefficient of variation in microbial network degree *C̄V̄* = 0.023). In contrast, the metabolic capacity of different genera varies by fourfold (**Fig. 2d**). For instance, species from the genus *Burkholderia* have >800 metabolites in their metabolic networks, while species from the genus Mycoplasma only have ∼200 metabolites in their metabolic networks. Matching our observations, the previously constructed genome-scale metabolic network for *Burkholderia cenocepacia* J2315 contains 834 metabolites^33^. By contrast, *Mycoplasma pneumoniae* was reported to have a small genome size, and its lifestyle is probably closer to obligate cellular parasites than to free-living organisms^34^. High metabolic similarity within genera and pronounced variation across genera suggest that metabolic capacities are evolutionarily conserved among closely related taxa^35^, while diversification across broader lineages might reflect specialization to distinct ecological niches^36^.

While the species degree distribution appears unimodal, the metabolite degrees display bimodality (**Fig. 2e**). This result indicates that many metabolites are specific to very few species, while many other metabolites are ubiquitous across the metabolic networks of almost all the 818 different species. Indeed, 57 (4.1%) of the 1390 metabolites (e.g., phlorizin and guaiacol) can only be processed by one or two species. This specificity may arise due to the unique enzymatic capabilities that those species possess, allowing them to process metabolites that others cannot. By contrast, 192 (13.8%) metabolites (e.g., cofactors, NADH, etc.) can be processed by more than 800 different species, possibly due to their essential role in microbial metabolism^37,38^.

We then compared the degree distributions of metabolites in different chemical classes (**Fig. 2f**). We found that peptidomimetics and some metal compounds can be processed by almost all species and may be necessary for microbes^39,40^. By contrast, azolidines, organic sulfuric acids/derivatives could only be processed by a few species, such as *Escherichia coli* and *Klebsiella pneumoniae*^41^. Furthermore, metabolites from many chemical classes, such as tannins, flavonoids, and keto acids/derivatives, exhibit a large variation in their degrees. For example, certain flavonoids (e.g., luteolin and quercetin) have complex structures that only a few microbial species can break down through specific enzymatic actions that other species cannot perform^42,43^. Other flavonoids (e.g., isoflavones) have simpler structures or more accessible chemical bonds, allowing a broader range of microbial species to metabolize them^44^.

The incidence matrix grouped by genera and classes of chemicals reveals similarity in preference for chemical classes within species of the same genus, and preference towards different chemical classes across genera (**Supplementary Figure 1**). For example, 91.8% ± 0.02% (mean ± std) of metabolites in the class of flavonoids exist in the metabolic networks of species from *Clostridium* and *Eubacterium*, significantly higher than the average presence of 44.5% ± 0.19% across all species (p-value = 4.42 × 10^!"#^, Mann-Whitney U Test). On the contrary, metabolites within the class of pyrimidine nucleotides exist in almost all species, likely due to the indispensability of pyrimidine nucleotides in synthesizing DNA and RNA^45,46^.

The preference of certain species for selected chemical classes is further confirmed by reorganizing the incidence matrix according to chemical and phylogenetic similarity (see Methods). Indeed, the reorganized incidence matrix similarly highlights the preference of some species for chemicals from particular classes (**Supplementary Figure 2**). Within this incidence matrix, the Jaccard similarity index for species within the same genus is 0.79 ± 0.08, which is significantly higher than that of 0.59 ± 0.09 for species from different genera (p-value < 10^!$##^, Mann-Whitney U Test; Rank-biserial correlation r_rb_=0.89 assuming from different genera and the same genus as group 1 and 2).

### Analysis of interactions between microbes and dietary compounds

Next, we narrowed the scope from the broader set of 1,390 metabolites to the 312 dietary compounds. We once again visualized the interactions between microbes and dietary compounds as a bipartite graph (**Fig. 3a**). We found that the incidence matrix of microbe–dietary compound interactions (NODF=0.870, **Fig. 3b**) is significantly more nested than the matrix of microbe– metabolite interactions (NODF=0.704, **Fig. 2b**), a result confirmed by randomly subsampling 312 metabolites out of 1,320 (p-value < 10^-4^; **Supplementary Figure 3**).

**Figure 3:**
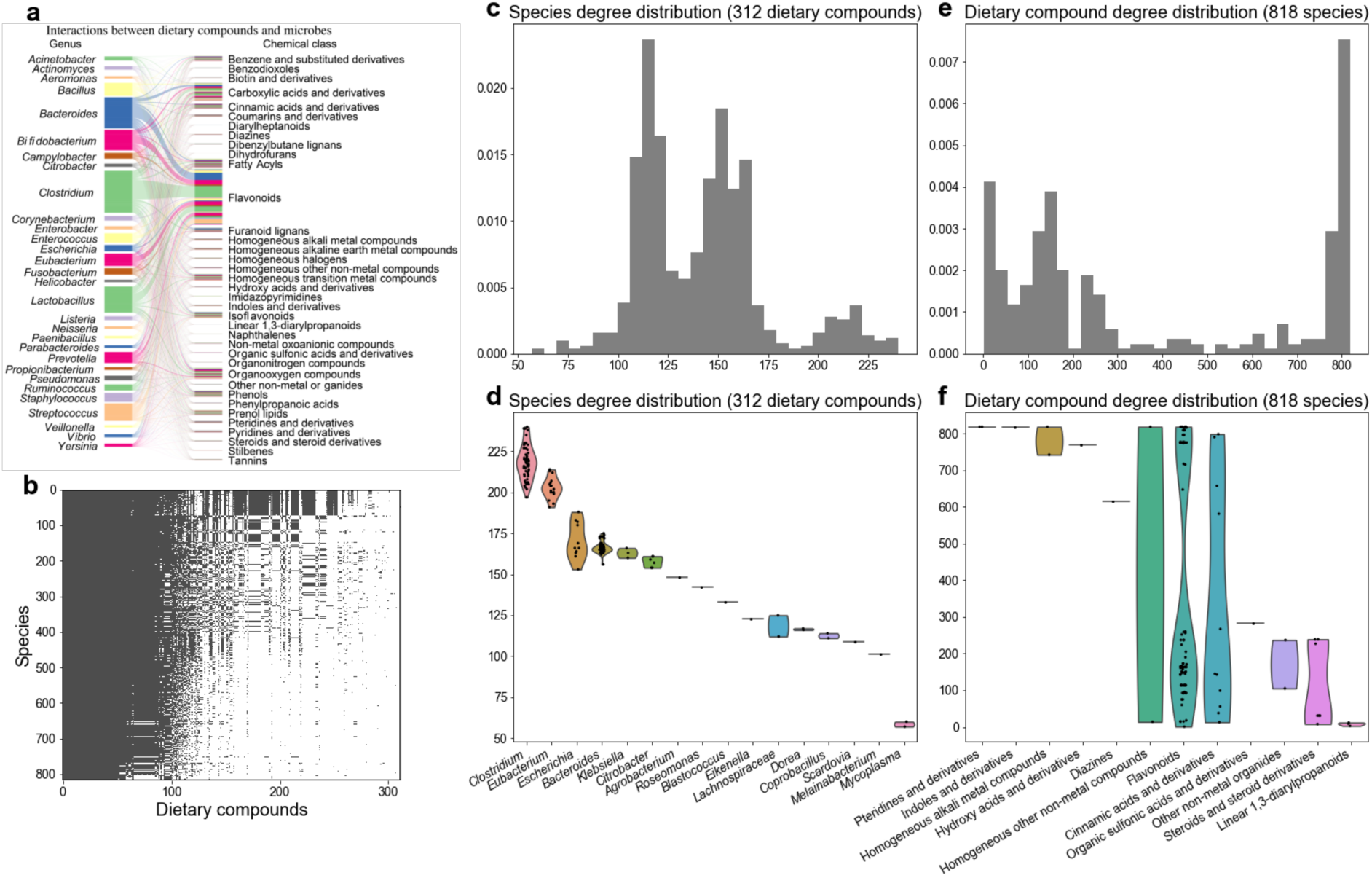
The inferred interactions between microbes and dietary compounds. **a,** The bipartite network of the interactions. To avoid visual overcrowding, we only displayed the top 30 genera and chemical classes with the largest counts. Each thin line on the left or right side corresponds to one species or dietary compound, respectively. Different species from the same genus and different dietary compounds from the same chemical class are grouped together without any gap in between. The curved line in the middle connects a species with a dietary compound when they interact. The color of nodes and edges is assigned based on the genus on the left. **b,** The incidence matrix of the interactions. The grey/white color for each element represents the presence/absence of a dietary compound in the metabolic network of a species. This matrix is organized using the Nestedness Temperature Calculator^71^ to emphasize its nested structure. **c,** The species degree distribution. The number of bins is 30. **d,** The species degree distribution for species from representative genera. **e,** The dietary compound degree distribution. The number of bins is 30. **f,** The dietary compound degree distribution from representative chemical classes.

Unlike **Fig. 2c**, the species degree distribution displays three main peaks: 110, 160, and 220 metabolites per species (**Fig. 3c**). This shows a higher degree of stratification in the number of dietary compounds metabolized by microbial genera, as further proven by **Fig. 3d**. For example, species from the genus *Clostridium* can utilize over 200 dietary compounds, which may explain the flexible metabolic capacity of *Clostridium* species reported in the literature^47^. In comparison, species from the genus *Mycoplasma* could use only approximately 50 dietary compounds, likely related to the small genome size for species from this genus such as *Mycoplasma pneumoniae*^34^.

Similar to the distribution of metabolite degrees, the network degree distribution of dietary compounds exhibits a predominant bimodality, with peaks around 0 and 800, as well as patterns of sub-peaks (**Fig. 3e**). Indeed, some dietary compounds, particularly those classified as linear 1,3-diarylpropanoids, can only be metabolized by a limited number of species, *Clostridium sp* and *Eubacterium ramulus* (**Fig. 3f**). In contrast, dietary compounds, like those in the chemical class of pteridines and derivatives, are present in the metabolic networks of most species. By focusing on dietary compounds with low degrees, which represent molecules restricted to the metabolic networks of a select few species, we can identify candidates with potential for targeting a narrow range of microbial species. Further examples and detailed analyses of these compounds are provided in the following sections.

By mapping microbial genera and chemical classes onto the incidence matrix (**Supplementary Figures 4–5**), we found distinct patterns in dietary compound prevalence across taxa. Flavonoids, the largest class in our dataset, are unevenly distributed, with a prevalence of 45.6% ± 19.4% (mean ± standard deviation) across species and are notably enriched in genera such as *Clostridium* and *Eubacterium*. Similarly, furanoid lignans show limited and variable presence (37.0% ± 25.3%), suggesting specialization. In contrast, carboxylic acids and derivatives exhibit high prevalence and a low standard deviation across all species (83.9% ± 4.1%), indicating their role as widely utilized compounds in core microbial metabolism.

### Relationship between genome size and network degree

We hypothesized that the variation in network degrees (or metabolic capacities) across species could be explained by their difference in genome sizes. To test this, we correlated the number of metabolites in the microbial genomes with their genome sizes, which also serves as evidence of annotation completeness. We visualize the statistical trend stratified by phyla (**Fig. 4a**). As expected, we found a significant correlation between them (Spearman Correlation Coefficient ρ = 0.59 with the p-value < 2.8 × 10^-75^). This agrees with our expectation that a microbial species with a bigger genome tends to contain more genes and can express more enzymes to catalyze chemical reactions (**Supplementary Figure 6**)^48–50^.

**Figure 4:**
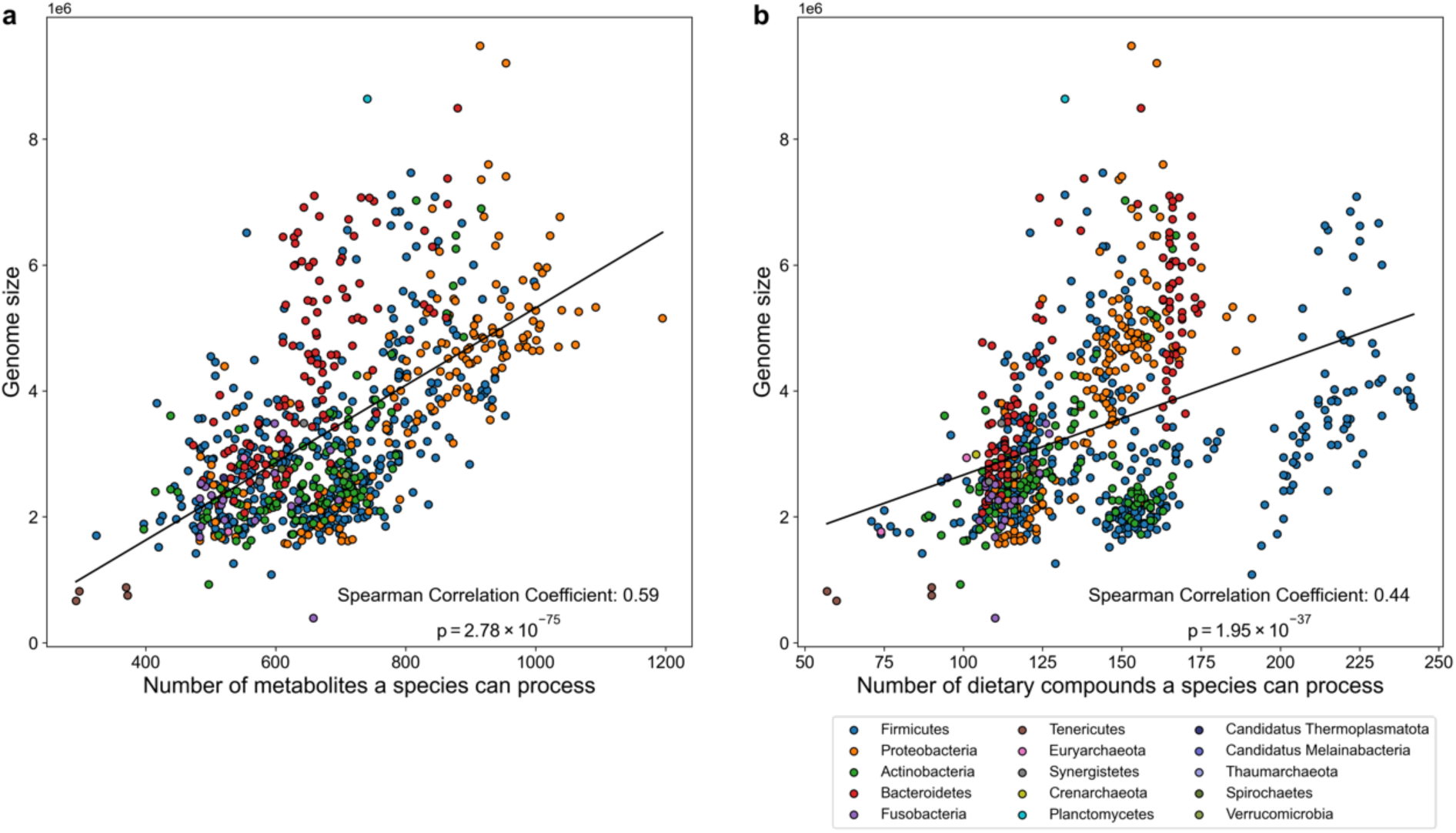
The positive correlation between species’ genome sizes and the number of metabolites or dietary compounds that species can process based on their genomes. In all figures, the dots are colored according to their phyla. **a,** The positive correlation between species’ genome sizes and the number of metabolites that species can process. **b,** The positive correlation between species’ genome sizes and the number of dietary compounds that species can process.

Similarly, we correlated the number of dietary compounds in the microbial genomes with their genome sizes, finding a significantly weaker correlation (Spearman Correlation Coefficient ρ = 0.44, p-value < 2.0 × 10^−37^ based on random samples of 312 metabolites in **Supplementary Figure 7**). This weaker correlation is likely due to a stronger stratification by phyla, where the number of metabolized dietary compounds appears less dependent on the genome size. For instance, a cluster of Firmicutes species exhibit a consistent range of 200 to 250 dietary compounds in their metabolic networks, despite their genome sizes varying from 10^’^ to 7.0 × 10^’^ bp (**Fig. 4b**). This pattern may reflect the high genomic and metabolic diversity among genera within the Firmicutes phylum, as previously reported^51^, which is greater than that observed in the more homogeneous Bacteroidetes phylum. We also expanded our analysis to the genus level, but the genus-level classification did not explain the separate cluster observed in our plot (**Supplementary Figure 8**).

For this cluster of Firmicutes, we investigated the dietary compounds with higher prevalence. Our enrichment analysis revealed a significant overrepresentation of flavonoids within this cluster: 61 out of the top 100 most over-represented dietary compounds in this cluster are flavonoids (p-value < 2.87 × 10^-8^, binomial test). This finding suggests a specialized metabolic adaptation to flavonoid processing within the phylum of Firmicutes, underscoring potential evolutionary and ecological implications for their role in the microbial community.

All microbial genomes exhibited high genome completeness (>92% for all), reflecting the use of high-quality reference genomes curated in the AGREDA database. To evaluate the potential confounding effect of genome completeness, we performed a multiple linear regression analysis in which the number of metabolites in the microbial genomes per species was modeled as a function of genome size while controlling for genome completeness. Genome size remained a highly significant predictor of the number of metabolites in the microbial genomes per species (β = 6.99 × 10^-5^, p = 6.97 × 10^-83^), indicating that genome completeness does not confound the observed relationship. A similar analysis for dietary compounds produced consistent results (β = 1.02 × 10^-5^, p = 5.11 × 10^-31^).

### Using the microbe-metabolite interactions to inform microbe-microbe interactions

Do metabolically similar microbes tend to (1) coexist due to habitat filtering^52,53^, where environmental conditions select for organisms with certain traits or adaptations that are most suited to a particular niche, or (2) compete, where stronger competitors in nutritional niches displace others who share similar metabolic traits^54,55^? To address this question, we quantified how metabolically similar microbes correlate with each other. Specifically, for any pair of microbial species, we measured the metabolic dissimilarity between the two species using the Jaccard distance between their abilities to process metabolites. Furthermore, we assessed their ecological interaction by calculating the Spearman correlation coefficient between the two species’ relative abundances across samples.

We utilized the MCTS dataset because it has time-series measurements spanning 17 days of gut microbial compositions for everyone. The high-frequency longitudinal nature of the data is instrumental in accurately quantifying dynamic microbe-microbe interactions on an individual basis. We found that the relative abundances of metabolically similar species tend to correlate positively (**Fig. 5a-d**; **Supplementary Figure 9**), supporting the hypothesis that habitat filtering facilitates the coexistence of metabolically similar species by exploiting different aspects of the same ecological niches. The overall weaker correlation when limited to dietary compounds is because metabolites collectively define more complete nutritional niches than dietary compounds, which represent only a subset of all metabolites (**Fig. 5e**; **Supplementary Figure 10**). Notably, while habitat filtering has been observed in previous studies for the gut microbiome through the analysis of the co-occurrence of microbial taxa^53^, our approach differs significantly. We have utilized within-individual time-series abundance correlations, rather than relying on microbial co-occurrence for cross-sectional data. This approach allows us to capture more dynamic interactions and temporal variations within everyone’s gut microbiome. Consequently, our method provides a more robust analysis, as it reveals consistent trends across individuals. Note that the dietary intake of the participants was not required for this analysis.

**Figure 5:**
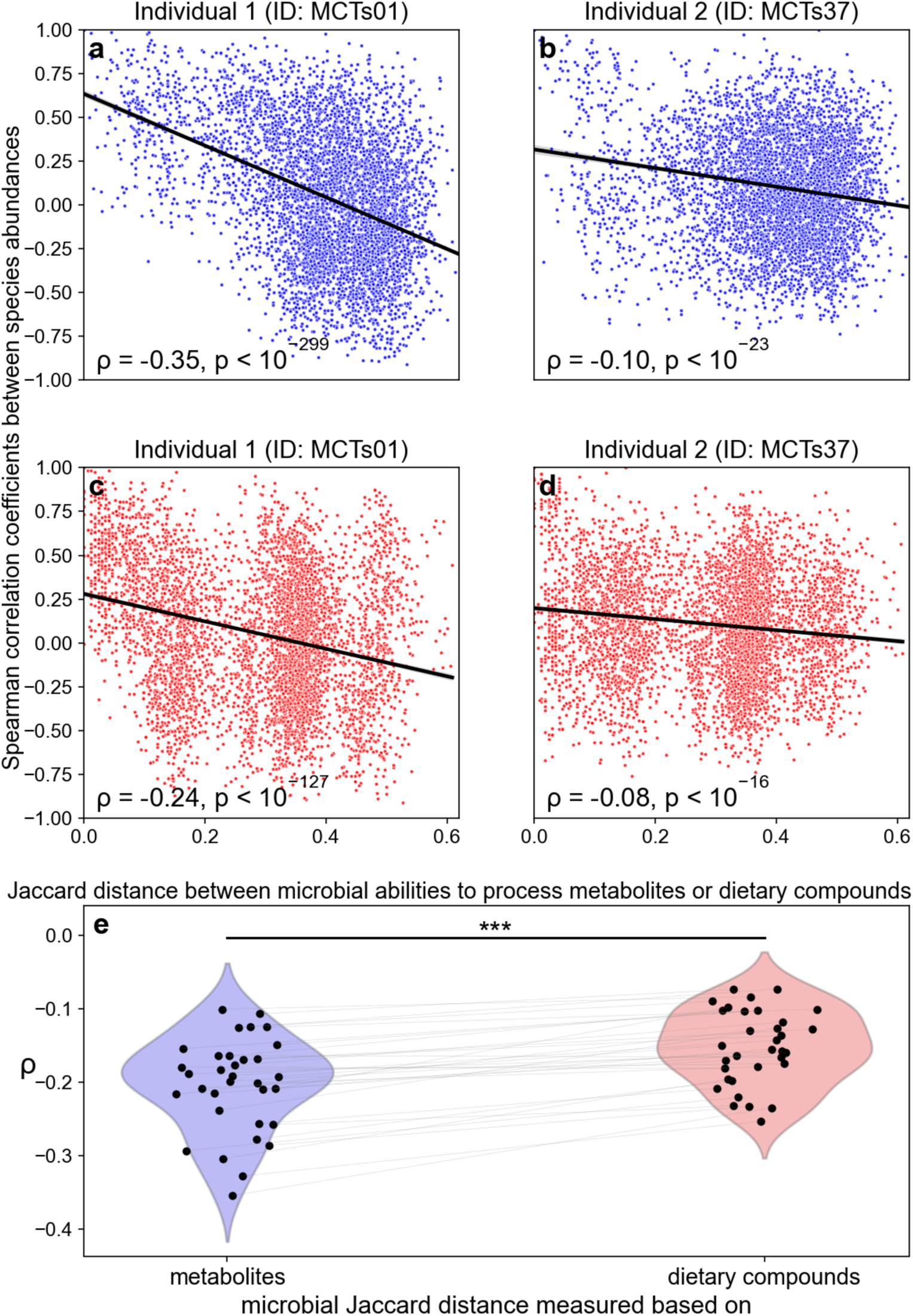
The relative abundances of metabolically similar species tend to correlate positively. Across all panels, the Spearman Correlation Coefficient between the relative abundances of two species (SCC_abun_) within the time-series gut microbial compositions for the same individual in MCTS^58^ is used to reflect the pairwise species-species ecological interaction. The Jaccard distance between two microbial species’ abilities to process metabolites (colored in blue) or dietary compounds (colored in red) is computed to reflect their similarity to process metabolites or dietary compounds. The Spearman Correlation Coefficient between SCC_abun_ and the Jaccard distance is denoted as :. The black lines are fitted linear regression results. **a,** For the individual with ID MCTs01, its SCC negatively correlates with the Jaccard distance between two microbial species’ abilities to process metabolites. **b,** For the individual with ID MCTs37, its SCC_abun_ negatively correlates with the Jaccard distance between two microbial species’ abilities to process metabolites. **c,** For the individual with ID MCTs01, its SCC_abun_ negatively correlates with the Jaccard distance between two microbial species’ abilities to process dietary compounds. **d,** For the individual with ID MCTs37, its SCC_abun_ negatively correlates with the Jaccard distance between two microbial species’ abilities to process dietary compounds. **e,** The correlation between the SCC_abun_ and Jaccard distance between two microbial species’ abilities to process metabolites is significantly smaller than the correlation between SCC_abun_ and the Jaccard distance between two microbial species’ abilities to process dietary compounds.

### The gut microbiome demonstrates large yet stable functional redundancy in its ability to process dietary compounds

Finally, we sought to leverage the interaction networks to explore the gut microbiome’s functional redundancy (FR)^31,56,57^ to process metabolites or dietary compounds: the extent of phylogenetically unrelated taxa to perform the same function. Following previous studies^31,56,57^, we first quantified the normalized gene-level functional redundancy 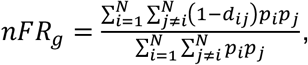 where *d_ij_* is the weighted Jaccard distance between the genomic contents of taxon-7 and taxon-*j* and *p_i_* is the relative abundance of taxon-7 (see Methods for the definition details). We extended the concept to define the normalized dietary-compound- and metabolite-level nFR by replacing the genomic contents with the genomic capability to process dietary compounds and metabolites, respectively.

Using the MCTS dataset, we quantified the three types of functional redundancies (nFR_d_, nFR_m_, and nFR_g_) for each time point across all individuals (**Supplementary Figure 11**). To assess overall trends, we quantified the mean level of nFR_d_, nFR_m_, and nFR_g_ for everyone by averaging across their respective time points. Our results revealed that, on average, nFR_d_ is higher than both nFR_m_ and nFR_g_ (**Fig. 6a-c**), likely due to the highly nested structure of the interaction network of microbes and dietary compounds. Such a nested network structure allows many microbes to share overlapping capacities for processing dietary compounds, resulting in high nFR_d_ values. This interpretation aligns with previous studies highlighting nestedness and functional redundancy in microbial metabolic networks^31^.

**Figure 6:**
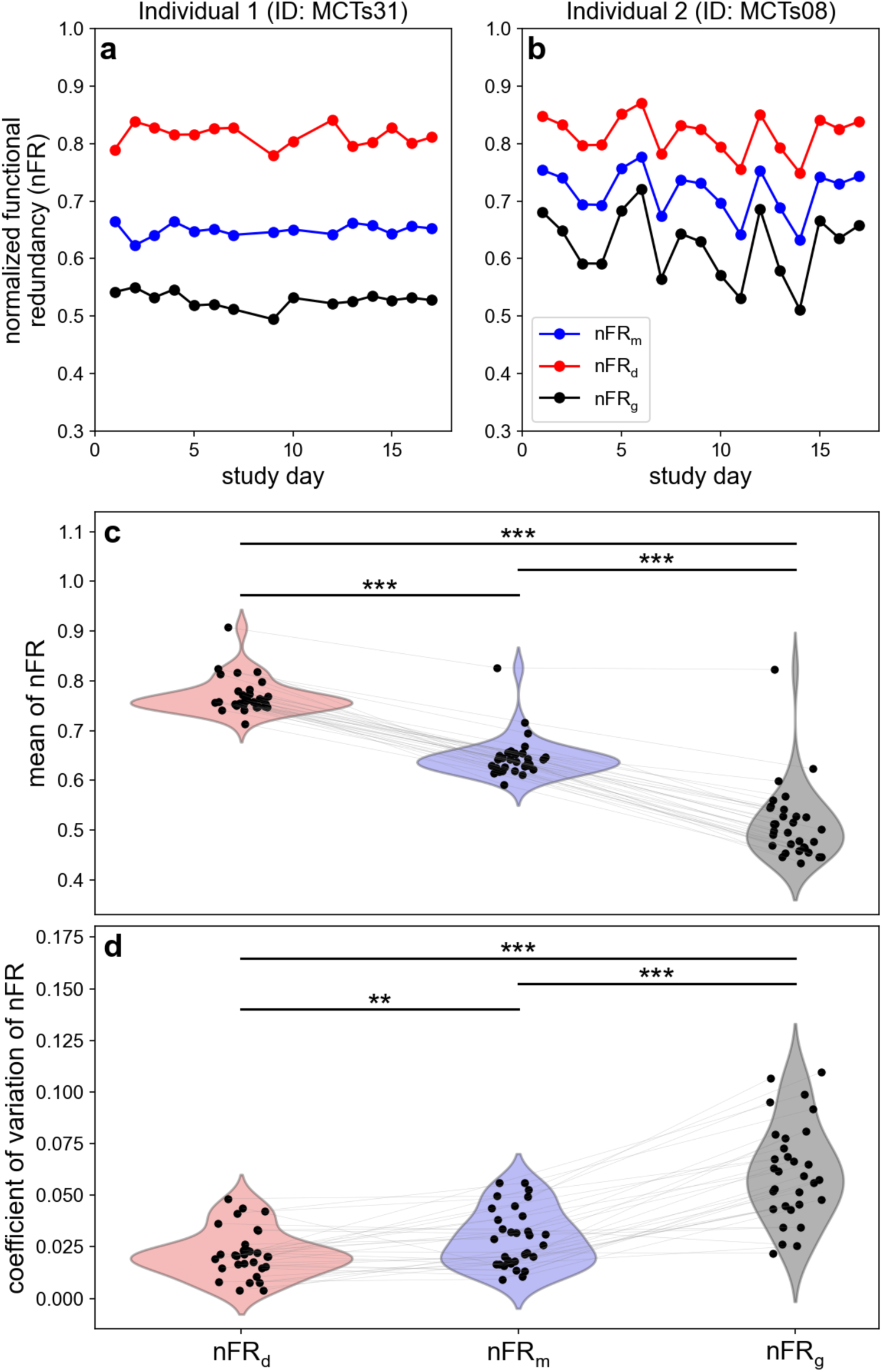
The normalized functional redundancy (FR) of the gut microbiome to process dietary compounds is higher and more stable than that of the gut microbiome to process metabolites or the gene-level functional redundancy. Here, nFR_d_, nFR_m_, and nFR_g_ refer to the normalized dietary-compound-, metabolite-, and gene-level FR, respectively. nFR_d_, nFR_m_, and nFR_g_ are calculated based on three networks of microbes interacting with metabolites, dietary compounds, and protein families respectively. **a,** The individual with ID MCTs31 has the most stable functional redundancies. **b,** The individual with ID MCTs08 has the most unstable functional redundancies. **c,** The mean of nFR_d_ is significantly larger than the mean of nFR_m_ and nFR_g_. **d,** The coefficient of variation (CV) of nFR_d_ is significantly smaller than the CV of nFR_m_ and nFR_g_.

We also compared the stability of functional redundancies using the coefficient of variation, revealing that nFR_d_ is more stable than nFR_m_ and nFR_g_ (**Fig. 6d**). Additionally, we quantified the null expectation of nFR_d_ based on 312 randomly selected metabolites and found that nFR_d_ is consistently higher than its null expectations (**Supplementary Figure 11**), suggesting that dietary compounds are uniquely different from random subsets of metabolites. These findings highlight the flexible yet robust nature of the gut microbiome’s ability to process dietary compounds.

### Specialized dietary compounds can alter the abundances of targeted species

To precisely target a narrow range of bacterial species through dietary intervention, it is crucial to identify the specialized dietary compounds that are exclusively linked with those species. To narrow down the list of microbial species to focus on, we only kept the microbial species in AGREDA that are present in the fecal samples of a real dataset, MCTS (microbiome diet study)^58^. To uncover these specialized dietary compounds, we focused on all dietary compounds with no more than ten species that contain them in their metabolic networks, identifying 49 specialized dietary compounds. We then selected all microbial species that possess at least one of these specialized dietary compounds in their metabolic networks, yielding a set of 63 species. The threshold of 10 species can be varied, and here we selected this threshold to balance interpretability and visual clarity, as lower thresholds (e.g., 5 species in **Supplementary Fig. 12a**) highlight only a few specialized compounds, while higher thresholds (e.g., 15 species in **Supplementary Fig. 12b**) result in network overcrowding. To gain further insights into the interactions between the 49 specialized dietary compounds and the 63 specialized species, we visualized their interactions as a network (**Fig. 7**). The complete list of the 49 specialized dietary compounds and their associated 63 microbial species is provided in **Supplementary Data 3**.

**Figure 7:**
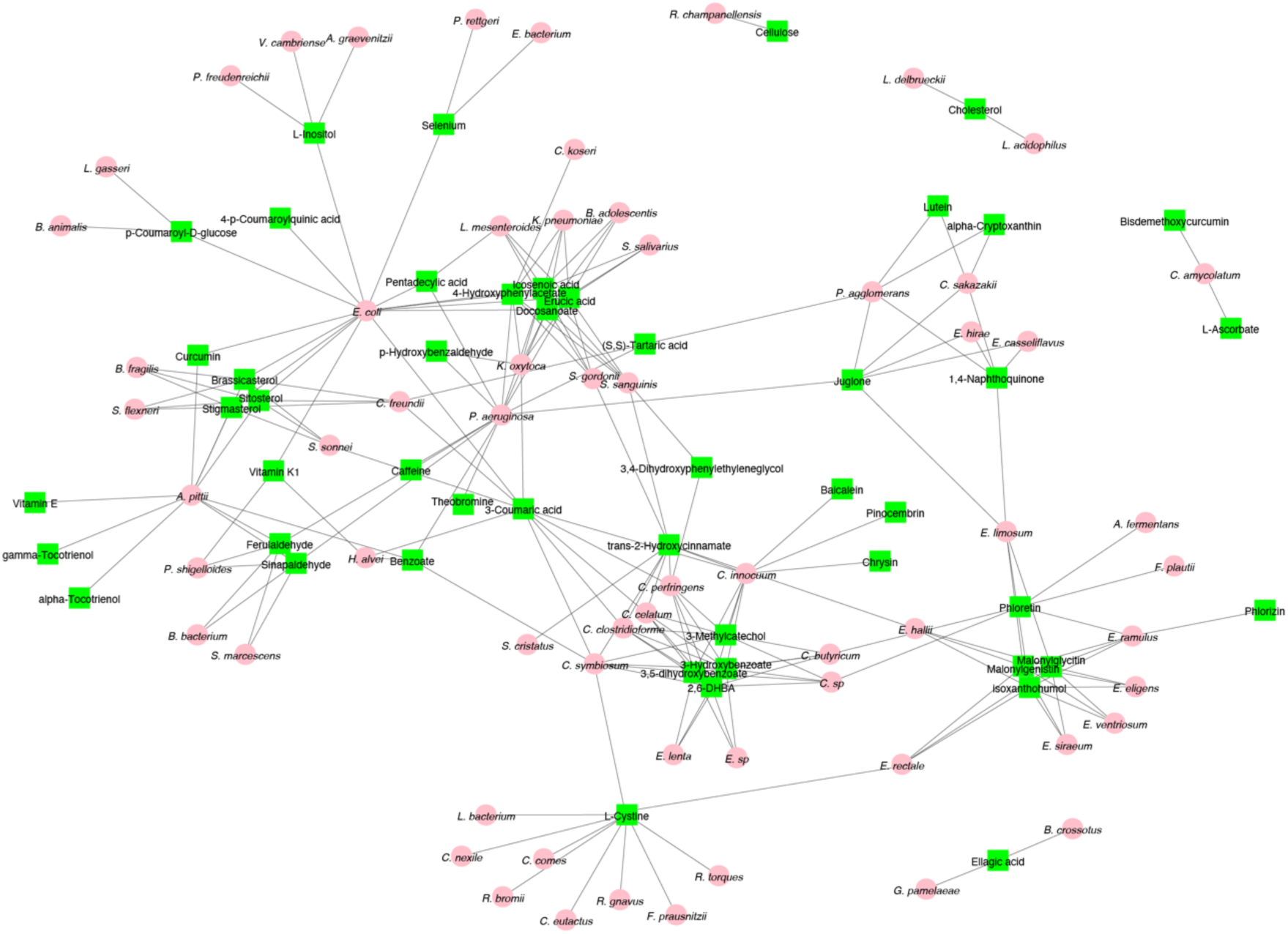
The interactions between specialized dietary compounds with linked bacteria. The specialized dietary compounds are present in no more than 10 microbial metabolic networks and their associated microbial species. The pink circles are microbial species, and the green squares are the dietary compounds. A black thin line connects a dietary compound with a microbial species when the dietary compound is present in the metabolic network of the species.

We hypothesize that one specialized dietary compound can be paired with its linked beneficial microbial species to design synbiotics (i.e., combinations of probiotics and prebiotics). For instance, let us consider the interaction between *Ruminococcus champanellensis (Rc)* and cellulose. *Rc* has been identified as a beneficial microbial species enriched in healthy individuals^59,60^, and the *Ruminococcus* genus has been shown to be less abundant in industrialized societies compared to ancient human, hunter-gatherer, and rural populations^60^. This reduction suggests a potential loss of beneficial gut functionality in contemporary populations. By using cellulose as a prebiotic to specifically support *Rc*, we hypothesize that the synbiotic combination could help restore the abundance of beneficial *Ruminococcus* species. Our hypothesis is validated by previous epidemiological studies showing that higher dietary intake of cellulose correlates with the increased abundance of *Ruminococcus*^60^.

We also investigated the possibility of leveraging the specialized interactions between *Eubacterium ramulus (Er)* and phlorizin to design synbiotics. *Er* is a quercetin-3-glucoside degrading anaerobic microorganism capable of degrading many flavonoids, including phlorizin^61^. *Er* converts these compounds into bioactive metabolites with antioxidant and anti-inflammatory properties that promote human health^61^. A previous human study demonstrated that the growth of *Er* can be stimulated by dietary flavonoids *in vivo*^62^, supporting the idea of *Er*-phlorizin synbiotics to boost the growth of *Er*.

## Discussion

We have systematically explored the interactions between microbes and metabolites, including dietary compounds based on inferred metabolic networks from well-annotated genomes of human gut microbes. While the present study did not generate new genome-scale reconstructions, it builds on the extensively validated AGREDA database to extract systems-level insights into gut microbial metabolism. Our primary objective here was to establish a network-level framework that integrates microbe–metabolite and microbe–dietary compound interactions to elucidate functional organization and inform the rational design of synbiotics. Specifically, we observed highly similar patterns of present metabolites and dietary compounds for species from the same genera and considerable variations in the presence of metabolites and dietary compounds within the metabolic networks for species across genera. In addition, we showed a positive correlation between the number of metabolites and dietary compounds in species’ metabolic networks and their genome sizes. We also explored the ecological relevance of these networks by quantifying the ecological interactions of metabolically similar species and the functional redundancy of the entire microbiome. Furthermore, we identified certain specialized dietary compounds that might be utilized and demonstrated their usefulness in modulating linked species. All insights gained from our study offer valuable information that can contribute to developing network-informed synbiotics.

Our finding that metabolically similar species co-occur supports the concept of habitat filtering, where the gut environment—shaped by host and dietary factors—selects for microorganisms with overlapping metabolic functions^52,53^. This process maintains functional redundancy within the microbiome, enabling diverse taxa to perform similar roles. Comparable patterns have been observed in non-industrialized populations, which often exhibit high microbial diversity despite relatively simple or seasonal diets^63^. Together, these observations suggest that selection acts primarily at the functional level.

The extension of microbial metabolism of dietary compounds through gap-filling and enzyme promiscuity methods^23,24^, such as those enabled by tools like RetroPath RL, has made the understanding of gut microbiota metabolism more complete. However, these approaches have limitations due to the existence of several equally plausible models, the inclusion of reactions lacking genomic evidence, and a deficit in experimental validation^64,65,50^. In addition, it is hard to distinguish whether a metabolite is consumed or produced only based on the enzymatic information. Their limitations underscore the need for improved computational models, integration of multi-omic data, and experimental validation. Together, these improvements are essential to fully elucidate the interactions between microbes, metabolites, and dietary compounds as well as their implications for human health.

It is important to acknowledge that our current results are constrained by the limited availability of high-quality, well-annotated microbial genomes. Although AGREDA provides one of the most comprehensive sets of curated gut microbial metabolic reconstructions to date, it still represents only a subset of the total diversity observed in metagenomic studies. Many rare, strain specific, or yet-uncultured taxa remain underrepresented, limiting the resolution of our inferred microbe–metabolite networks. In the future, with more sequencing of microbial genomes, we can expand the number of included species in our analysis. In this fashion, as the knowledge of gene annotation improves, it will help uncover more interactions between microbes and metabolites. Although our study focused solely on the nutrient constituents of foods (i.e., dietary compounds), it is worth noting that food intervention is a more feasible approach than direct intervention on dietary compounds in real-life scenarios. To accurately figure out which food shall be used to target certain microbial species, the comprehensive and accurate documentation of dietary compounds within each food is necessary.

## Methods

### Genome-scale metabolic models (GEMs) of microbes

The metabolic networks used were retrieved from the GEMs documented in AGREDA^23,24^, which provides information about annotated enzymes and catalyzed chemical reactions for 818 gut microbial species. The link to the deposited GitHub repository can be found here: https://github.com/francescobalzerani/AGREDA_1.1. All chemical reactions in its metabolic model were pulled out for each microbial species, and then all metabolites that exist in any chemical reactions were captured. This process of extracting metabolites was repeated for all microbial species. Eventually, all interactions between microbial species and metabolites were summarized in a binary network or an incidence matrix.

### Dataset

The dataset we leveraged to test the idea of targeted microbiota modulation comes from MCTS (MiCrobiome dieT study)^58^, which collected the paired fecal samples and dietary intake for 34 participants over 17 consecutive days. The microbial composition of fecal samples was measured by 16S sequencing. The intake of all foods, drinks, and supplements in the past 24 hours was captured by ASA24 (Automated Self-Administered 24-Hour) based on participants’ recall. The raw and processed data can be found in the supplementary information of the paper that investigated the diet-microbiome associations in MCTS^58^.

### Chemical structure and property

AGREDA provides chemical identifiers such as ModelSEED compound ID^66^, InChIKey (International Chemical Identifier hash Key)^67^, SMILES (Simplified Molecular Input Line Entry System)^68^, and KEGG KOs (Kyoto Encyclopedia of Genes and Genomes Orthology)^69^. The structural similarity between any two chemicals was computed based on the similarity between the Morgan fingerprints of the two SMILES strings^70^. Specifically, for each atom within a chemical, a series of circular expansions was performed to obtain unique substructures. The maximal radius we used in the expansion is 3. Then, a binary representation with 8096 bits was used to reflect if each unique substructure exists in the chemical. Eventually, the Jaccard similarity index between two bitstrings was calculated to measure their structural similarity. For each chemical, we also used its InChIKey to find its chemical classes in ClassyFire^30^, a web-based application for the automated structural classification of chemical entities.

### Clustering by metabolite chemical similarity and microbial phylogenetic similarity

We restructured the incidence matrix by clustering metabolites and species according to chemical structure and phylogenetic similarity. We utilized the SMILES representation of chemicals, which represents structure by capturing the chemical elements and bonds between elements^68^. To compute the similarity in chemical structure, we employed the Morgan fingerprint, which is derived from a series of circular substructures around each atom and then generated by a binary representation for a given chemical compound based on its molecular structure within each circular substructure^70^. To measure the microbial similarity, we leveraged the phylogenetic similarity.

### Nestedness

The nested property of the incidence matrix was visualized by organizing it via the Nestedness Temperature Calculator^71^. The degree of nestedness was measured by NODF (Nestedness metric based on the overlap and decreasing fill)^72^.

### Functional redundancy

We directly used the normalized gene-level functional redundancy (*nFR_g_*) defined in previous studies: 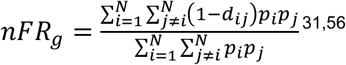, where *d_ij_* is the functional distance between taxon-*i* and taxon-*j* measured by the weighted Jaccard distance between their genomic contents (i.e., genome-annotated protein families such as KOs for each taxon) and *p_i_* denotes the relative abundance of taxon-7 within the microbial community. Similar to this definition of *nFR_g_*, we extend the concept to define *nFR_d_* and *nFR_m_* by replacing the genomic contents with the genomic capability to process dietary compounds and metabolites, respectively.

### Statistics

To calculate correlation throughout the study, we used Spearman’s correlation coefficient. Wherever we used *P* values, we explained in the main text how we calculated the standard statistical tests or the null distributions from scratch. All statistical tests were performed using standard numerical and scientific computing libraries (Numpy^73^, Scipy^74^, Pandas^75^, Matplotlib^76^, and Seaborn^77^) in Python (version 3.7.3) and Jupyter Notebook (version 6.1).

## Data and code availability

All metabolic networks in AGREDA can be found here: https://github.com/francesco-balzerani/AGREDA_1.1. Within this repository, the genome-scale metabolic networks of all microbial species in AGREDA are located in the folder: AGREDA_1.1/02_ManualRevisionAndReconstruction/output/outputReconstruction/singleSpecie s/. All data related to MCTS can be found in the following repository: https://github.com/knightslab/dietstudy_analyses. In this repository, microbial composition data for all samples can be found in this file: dietstudy_analyses/data/microbiome/processed_sample/taxonomy_counts_s.txt.

## Author contributions

T.W., Y.-Y.L., and G.M. designed the project. T.W. performed all the numerical calculations and data analysis. T.W. processed the real data. All authors analyzed the results. T.W., Y.-Y.L., and G.M. wrote the paper. All authors edited and approved the paper.

**Correspondence.** Correspondence and requests for materials should be addressed to Y.-Y. L. and G.M.

## Declarations

**Ethics approval and consent to participate.** Not applicable

**Consent for publication.** Not applicable

