## Supplemental Information for "Revealing Interactions between Microbes, Metabolites, and Dietary Compounds using Genome-scale Analysis"

**Supplementary Figures**

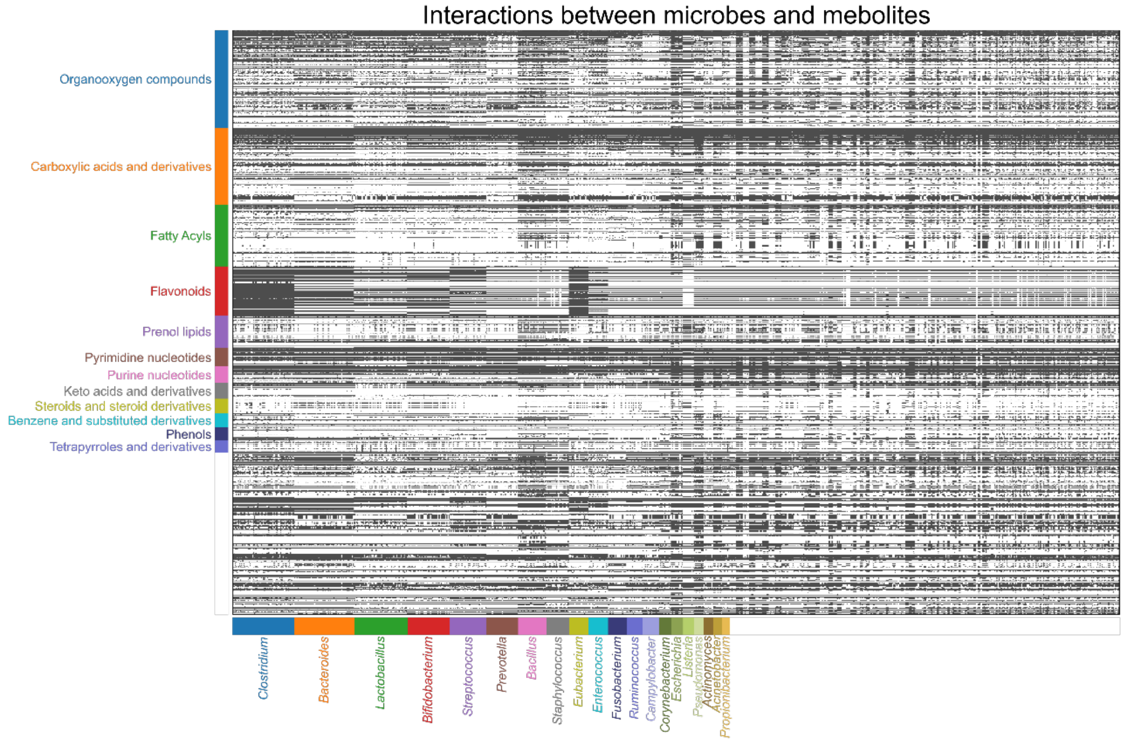


**Supplementary Figure 1: The incidence matrix of inferred interactions between microbes and metabolites**. The grey/white color for each element represents the presence/absence of a metabolite in the metabolic network of a species. From top to bottom, the matrix is organized following the decreasing counts of metabolites in all chemical classes. The names of the top 20 chemical classes with the most counts are denoted. From left to right, the matrix is organized following the decreasing counts of species in all genera. The names of the top 20 genera with the most counts are denoted.

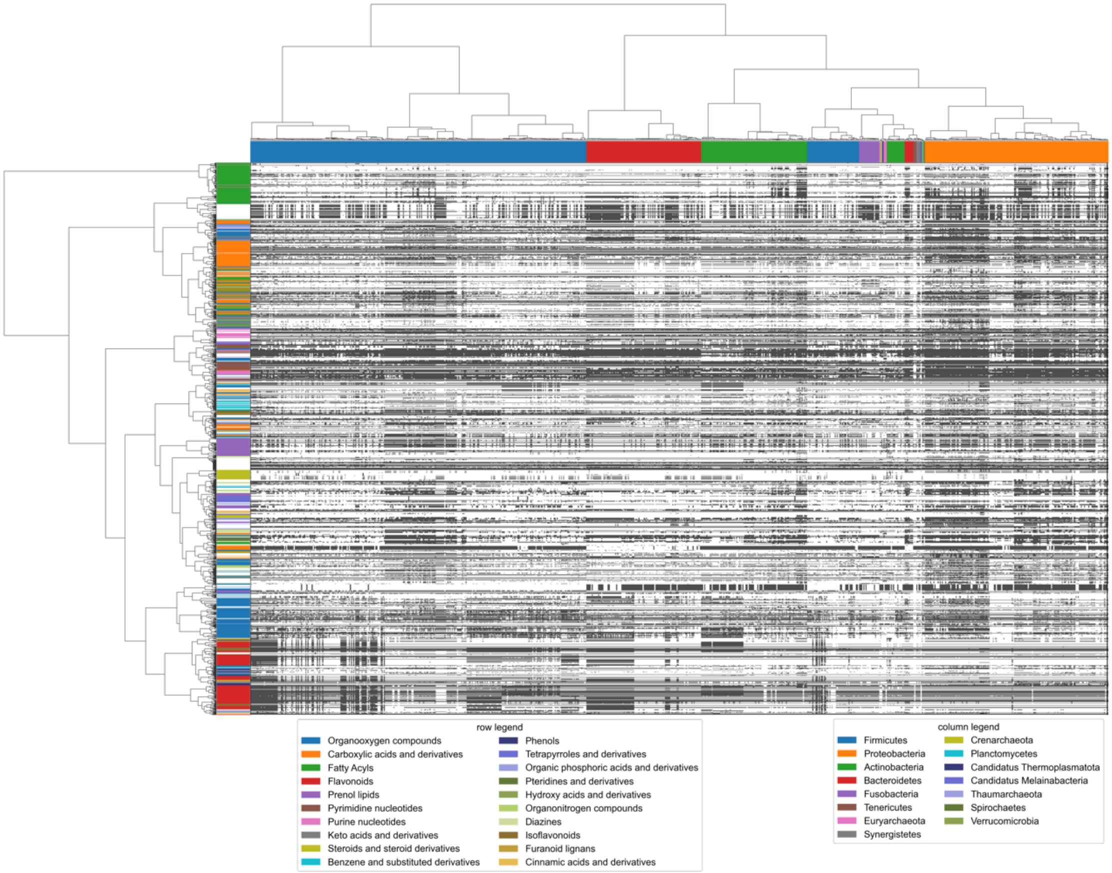


**Supplementary Figure 2: The incidence matrix of inferred interactions between microbes and metabolites**. The grey/white color for each element represents the presence/absence of a metabolite in the metabolic network of a species. The rows and columns of the heatmap are hierarchically clustered based on the structural similarity of Morgan fingerprints and phylogenetic similarity respectively. The color bar on the left denotes the top 20 chemical classes with the most counts of metabolites. The color bar on the top denotes the order-level taxonomy of microbial species.

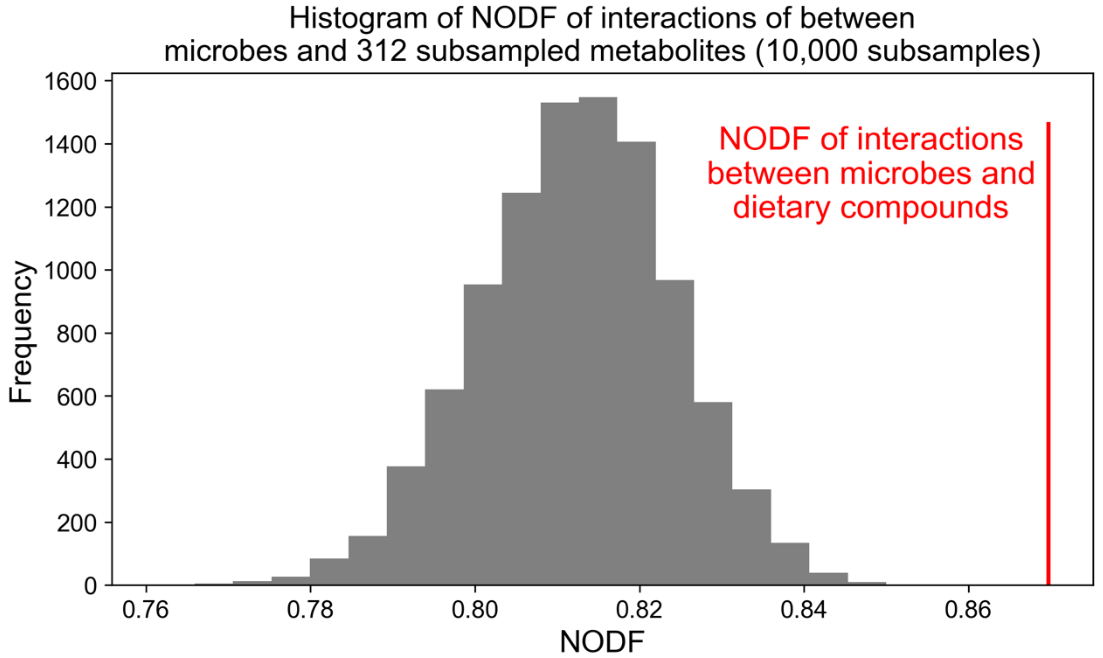


**Supplementary Figure 3: The interactions between microbes and dietary compounds are significantly more nested than the interactions between microbes and subsampled metabolites**. Out of 1390 metabolites, a random set of 312 metabolites is subsampled (same as the number of dietary compounds), then the nestedness of the subsampled matrix is evaluated via NODF. Such a subsampling procedure and calculation of NODF is performed 10,000 times to generate the histogram.

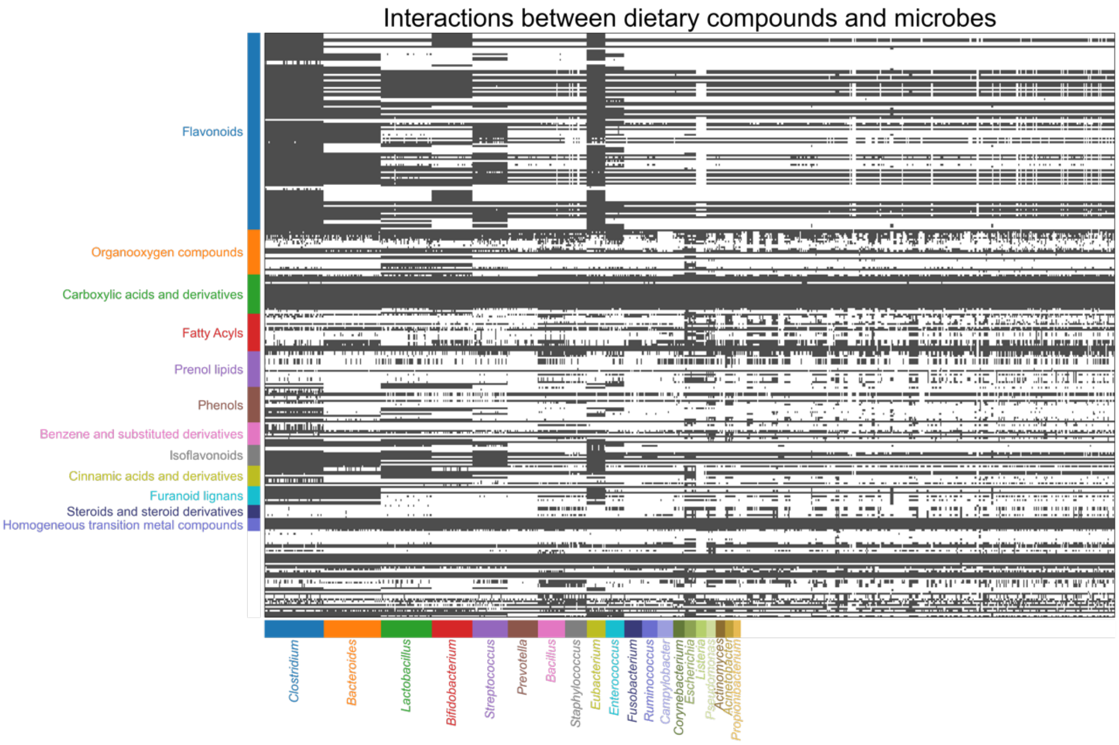


**Supplementary Figure 4: The incidence matrix of inferred interactions between microbes and dietary compounds**. The grey/white color for each element represents the presence/absence of a dietary compound in the metabolic network of a species. From top to bottom, the matrix is organized following the decreasing counts of dietary compounds in all chemical classes. The names of the top 20 chemical classes with the most counts are denoted. From left to right, the matrix is organized following the decreasing counts of species in all genera. The names of the top 20 genera with the most counts are denoted. The heatmap is hierarchically clustered.

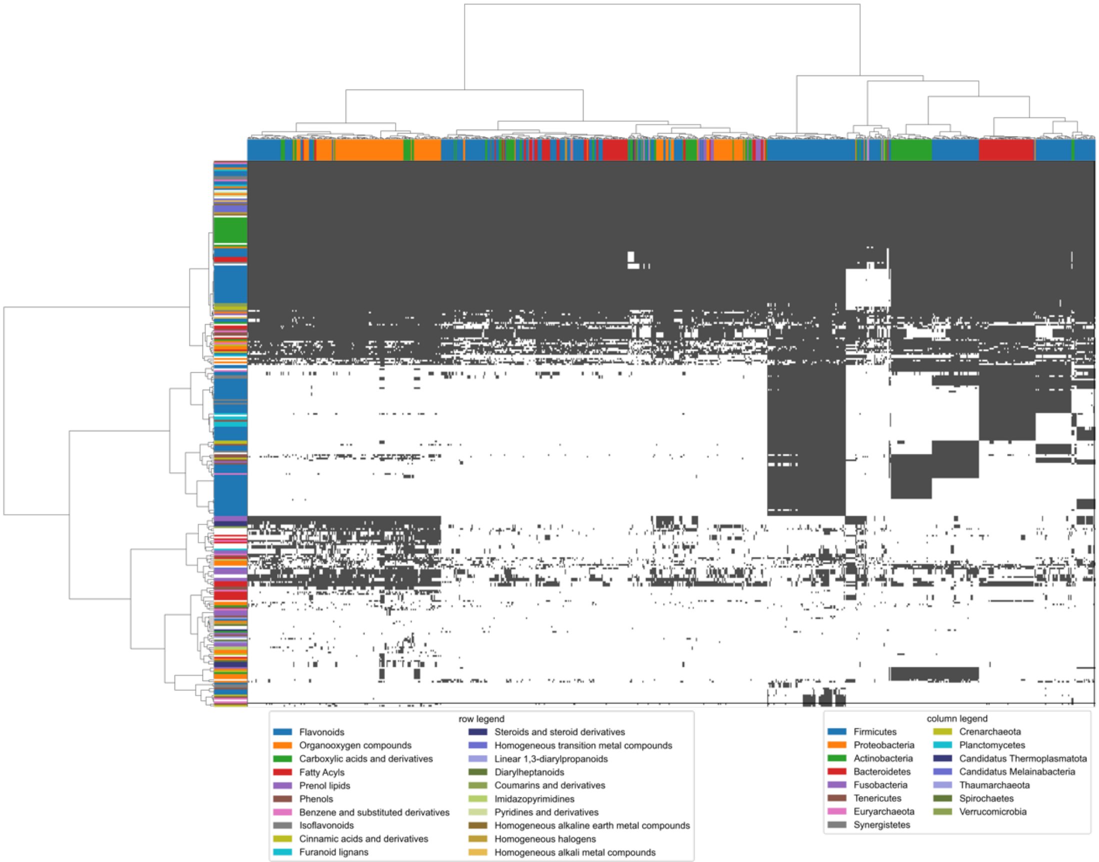


**Supplementary Figure 5: The incidence matrix of inferred interactions between microbes and dietary compounds**. The grey/white color for each element represents the presence/absence of a dietary compound in the metabolic network of a species. The rows and columns of the heatmap are hierarchically clustered based on the structural similarity of Morgan fingerprints and phylogenetic similarity respectively. The color bar on the left denotes the top 20 chemical classes with the most counts of dietary compounds. The color bar on the top denotes the order-level taxonomy of microbial species.

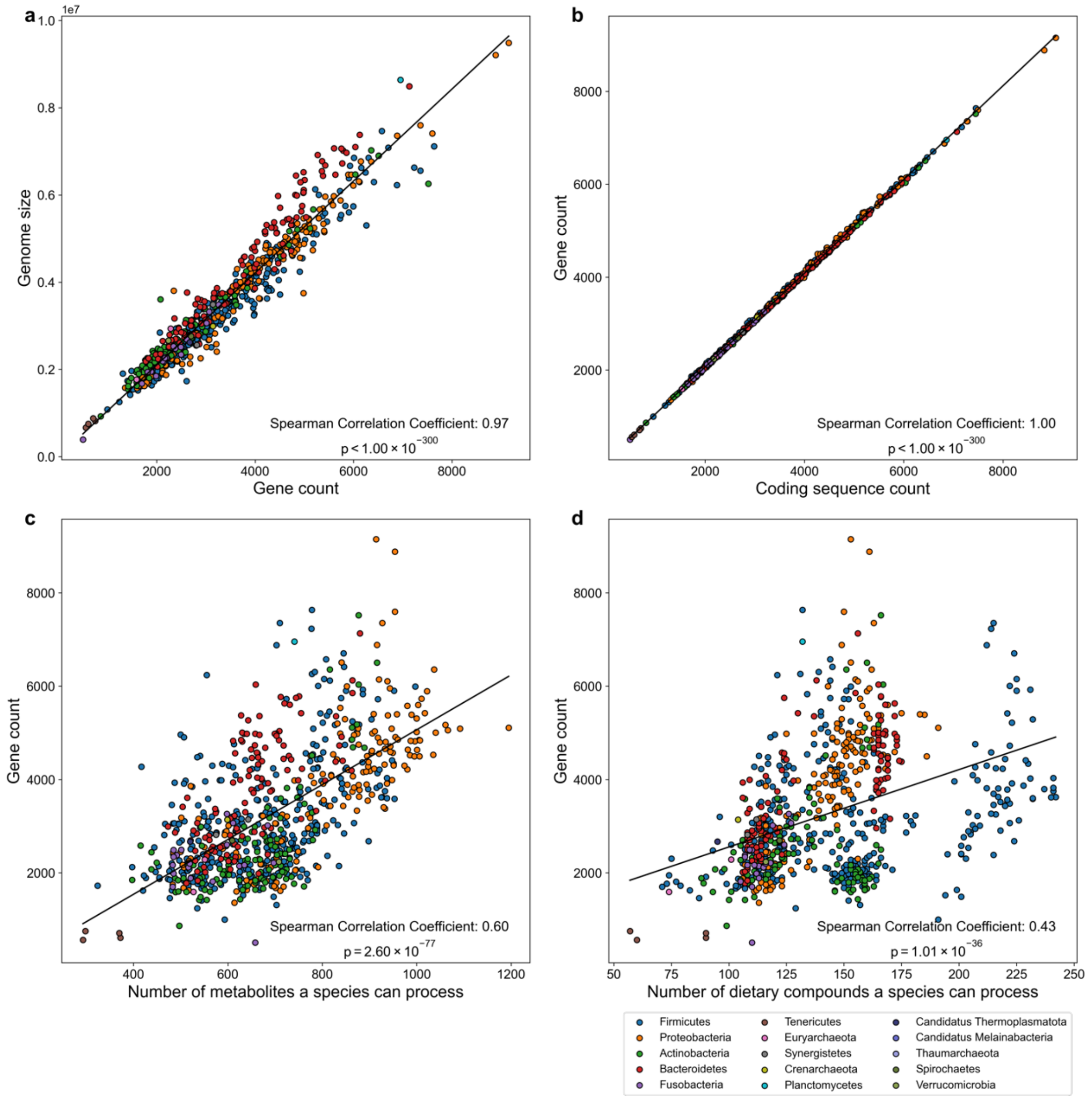


**Supplementary Figure 6: The positive correlation between species' genome sizes, species' gene count, species' coding sequence count, and the number of metabolites or dietary compounds that species can process based on their genomes**. In all figures, the dots are colored according to the order level taxonomy. **a,** The positive correlation between species' genome sizes and gene counts. **b,** The positive correlation between species' genome sizes and coding sequence counts. **c,** The positive correlation between species' gene counts and the number of metabolites that species can process. **d,** The positive correlation between species' gene counts and the dietary compounds that species can process.

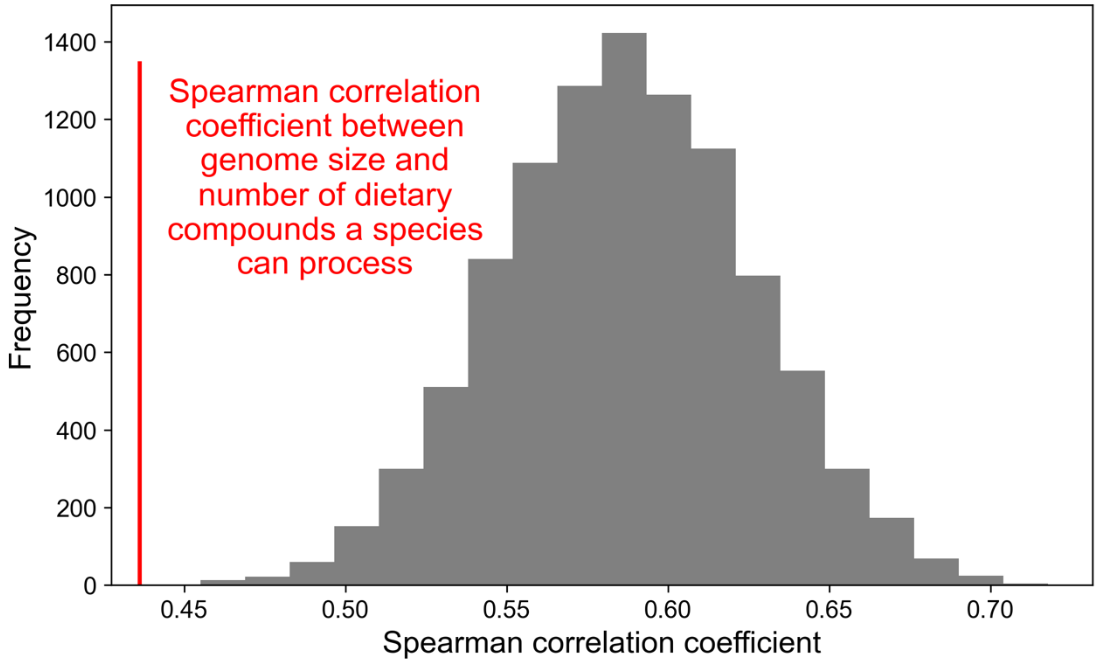


**Supplementary Figure 7: The Spearman correlation between the number of dietary compounds that can be processed by a species and the genome size of the species is significantly weaker than the Spearman correlation between the number of metabolites and genome size**. Out of 1390 metabolites, a random set of 312 metabolites is subsampled (same as the number of dietary compounds), then the Spearman correlation between the number of sampled metabolites and genome size is evaluated. Such subsampling procedure and calculation of correlation are performed 10,000 times to generate the histogram.

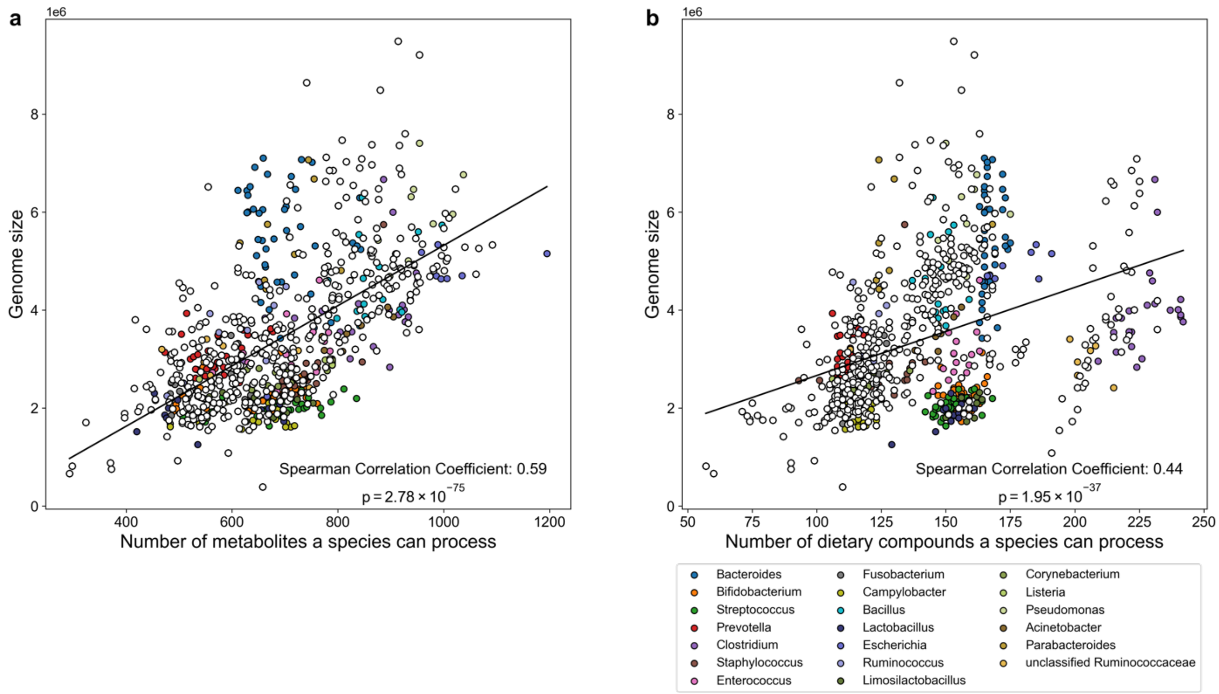


**Supplementary Figure 8: The positive correlation between species' genome sizes and the number of metabolites or dietary compounds that species can process based on their genomes**. In all figures, the dots are colored according to their genera. **a,** The positive correlation between species' genome sizes and the number of metabolites that species can process. **b,** The positive correlation between species' genome sizes and the number of dietary compounds that species can process.

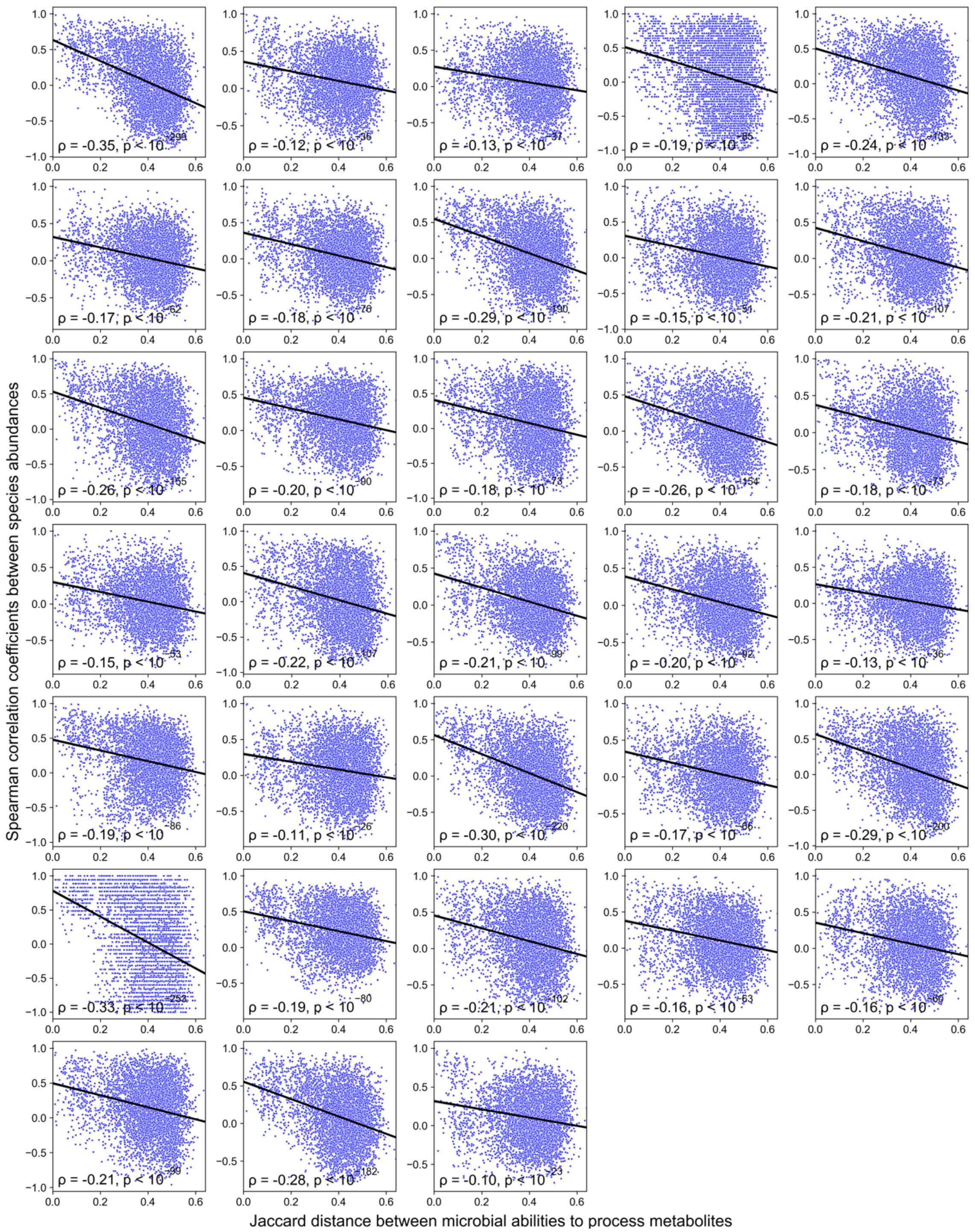


**Supplementary Figure 9: The relative abundances of metabolically similar species tend to correlate positively**. For each subplot, its data is derived from one individual in MCTS[58]. Spearman Correlation Coefficient $\rho$ is shown. The black lines are fitted linear regression results.

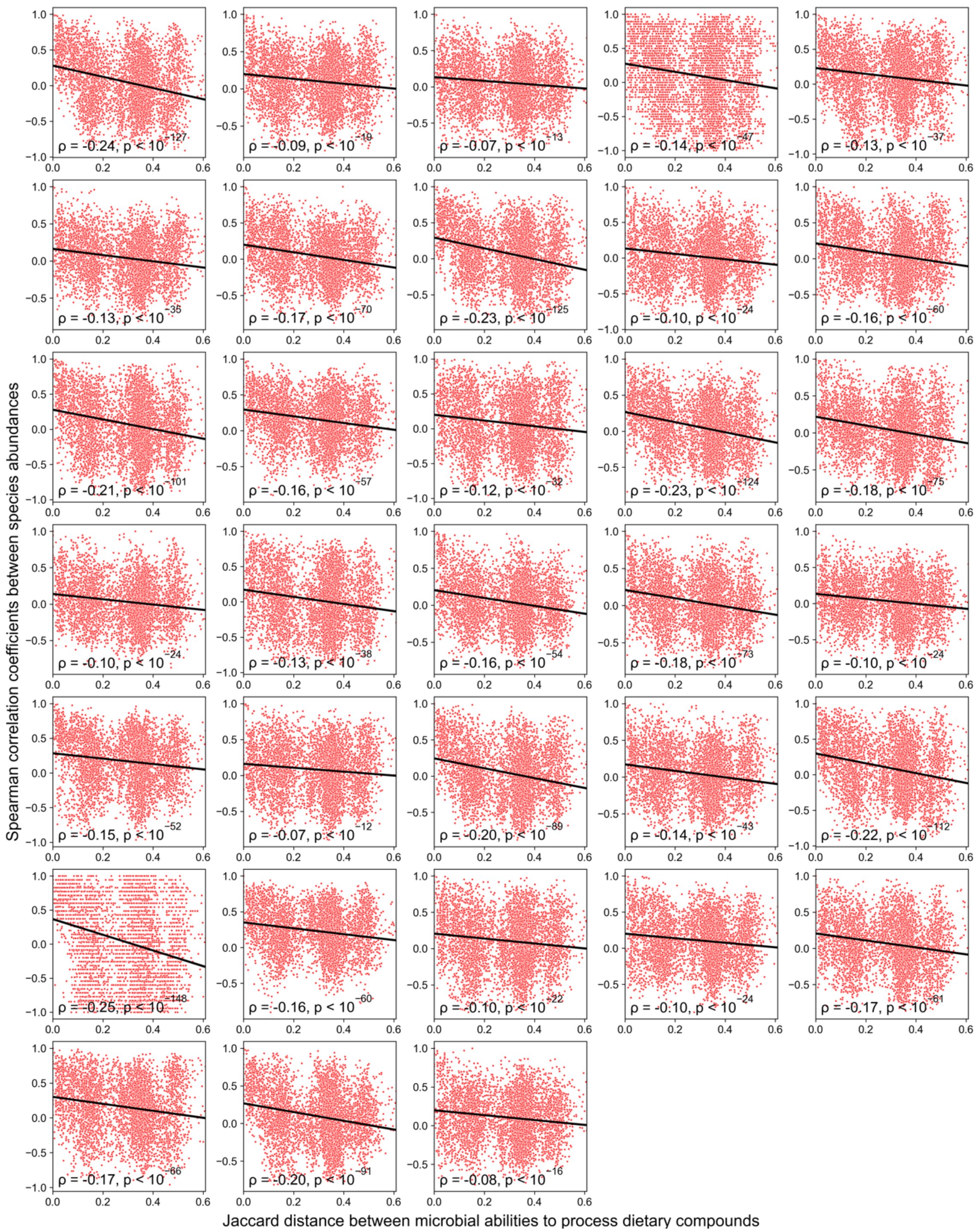


**Supplementary Figure 10: The relative abundances of species that share similar abilities to process dietary compounds tend to correlate positively**. For each subplot, its data is derived from one individual in MCTS[58]. Spearman Correlation Coefficient $\rho$ is shown. The black lines are fitted linear regression results.

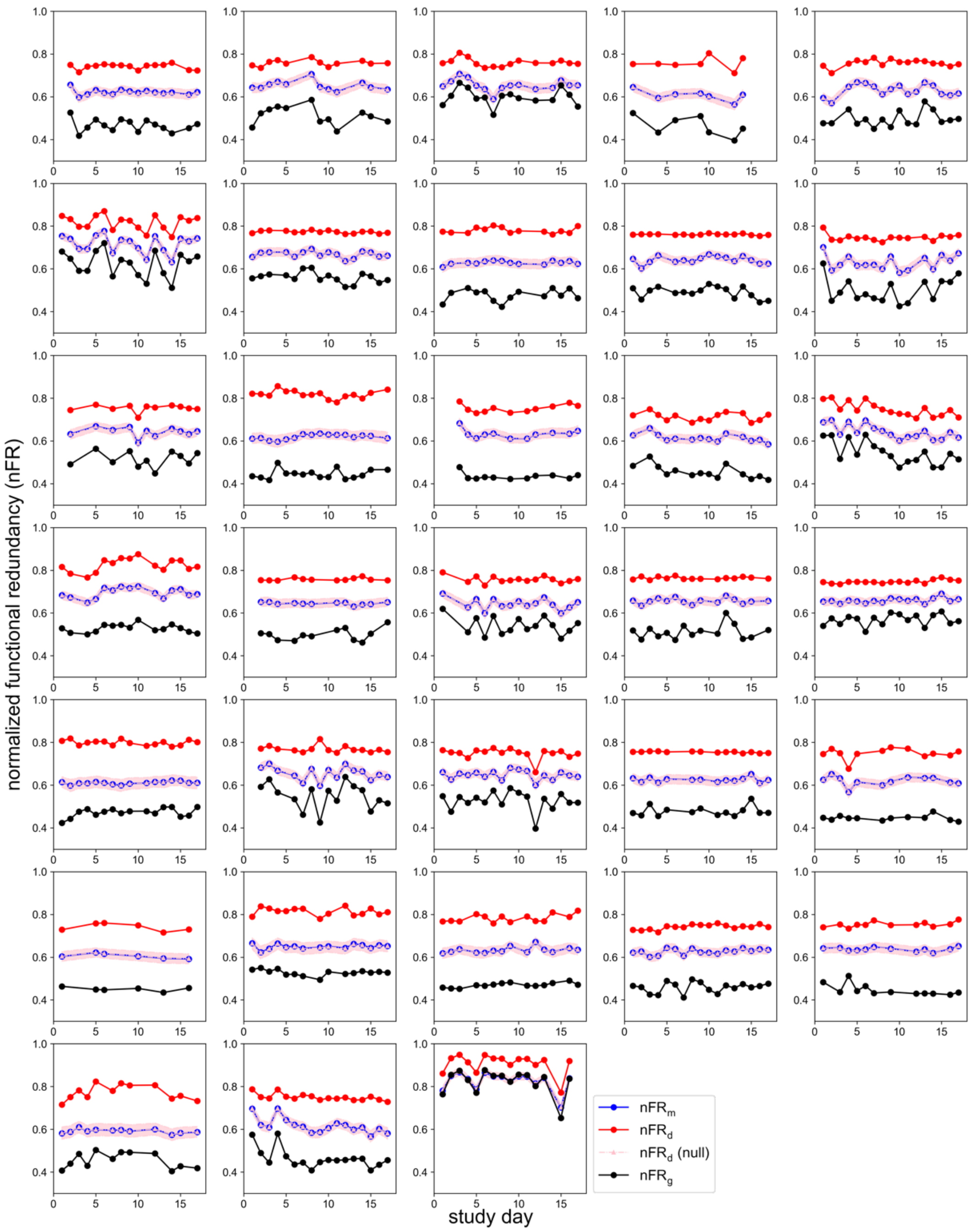


**Supplementary Figure 11: The normalized functional redundancy of the gut microbiome to process dietary compounds is higher and more stable than that of the gut microbiome to process metabolites or the gene-level functional redundancy**. Here, $nFR_d$, $nFR_m$, and $nFR_g$ refer to the normalized dietary-compound-, metabolite-, and gene-level FR, respectively. For each subplot, its data is derived from one individual in MCTS[58]. Out of 1390 metabolites, a random set of 312 metabolites is subsampled (same as the number of dietary compounds), then the null expectation of $nFR_d$ is calculated and denoted as $nFR_d$ (null). Such a subsampling procedure and calculation is performed 10,000 times to generate the mean (pink dashed line) and standard deviation (pink transparent area).

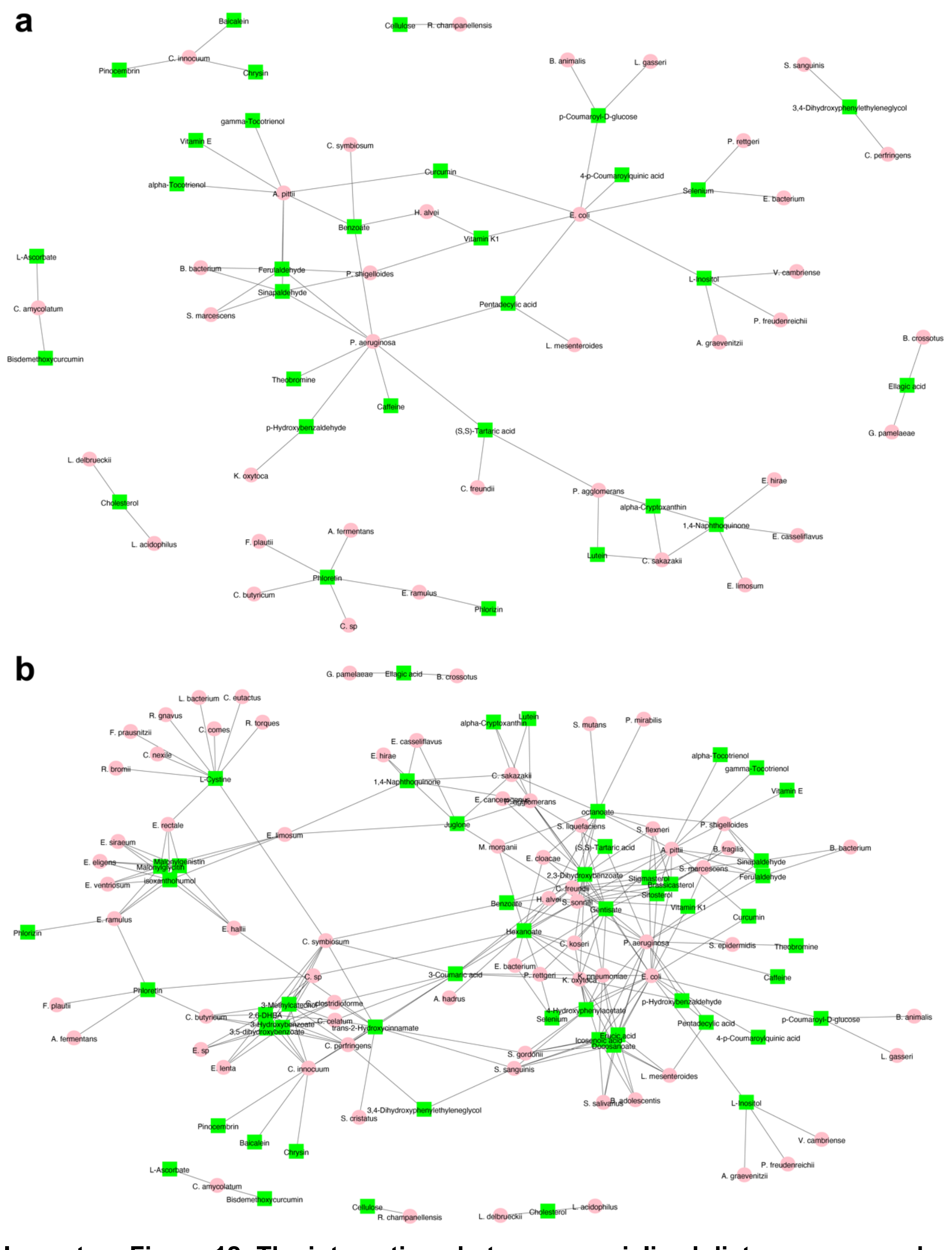

**Supplementary Figure 12: The interactions between specialized dietary compounds with linked bacteria**. The pink circles are microbial species, and the green squares are the dietary compounds. A black thin line connects a dietary compound with a microbial species when the dietary compound is present in the metabolic network of the species. **a,** The specialized dietary compounds are present in no more than 5 microbial metabolic networks and their associated microbial species. **b,** The specialized dietary compounds are present in no more than 15 microbial metabolic networks and their associated microbial species.
